# Calcium-Induced Differentiation in Normal Human Colonoid Cultures: Cell-cell / cell-matrix adhesion, barrier formation and tissue integrity

**DOI:** 10.1101/505016

**Authors:** Durga Attili, Shannon D. McClintock, Areeba H. Rizvi, Shailja Pandya, Humza Rehman, Daniyal M Nadeem, Aliah Richter, Dafydd Thomas, Michael K. Dame, D. Kim Turgeon, James Varani, Muhammad Nadeem Aslam

## Abstract

Colonoid cultures were established from histologically-normal human colon tissue and maintained in a low-calcium (0.25 mM) medium or in medium supplemented with an amount of calcium (1.5 - 3.0 mM) that was shown in a previous study to induce differentiation in colonoids derived from large adenomas. Calcium alone was compared to Aquamin, a multi-mineral natural product that contains magnesium and detectable levels of 72 additional trace elements in addition to calcium. Unlike the previously-studied tumor-derived colonoids (which remained un-differentiated in the absence of calcium-supplementation), normal tissue colonoids underwent differentiation as indicated by gross and microscopic appearance, a low proliferative index and high-level expression of cytokeratin 20 (CK20) in the absence of intervention. Only modest additional changes were seen in these parameters with either calcium alone or Aquamin (providing up to 3.0 mM calcium). In spite of this, proteomic analysis and immunohistochemistry revealed that both interventions induced strong up-regulation of proteins that promote cell-cell and cell-matrix adhesive functions, barrier formation and tissue integrity. Transmission electron microscopy revealed an increase in desmosomes in response to intervention. These findings demonstrate that histologically normal human colonoids can undergo differentiation in the presence of a low ambient calcium concentration. However, higher calcium levels can induce elaboration of proteins that promote cell-cell and cell-matrix adhesion. These changes could lead to improved barrier function and improved colon tissue health.

## INTRODUCTION

Epidemiological studies have demonstrated that calcium intake and colon polyp formation / colon cancer are inversely related (1-7). In spite of this, interventional trials using calcium supplementation to reduce polyp formation have been only modestly effective. Some chemoprevention trials have shown a reduction in polyp incidence (8,9) but others have failed to find significant benefit (10,11). One chemoprevention study actually found an increased risk of developing colon polyps with the sessile-serrated phenotype – a pathological presentation associated with increased risk of colon cancer - with calcium supplementation (12). The same interventional trials have also demonstrated only modest (though in some cases, statistically significant) effect on biomarkers of growth and differentiation in histologically-normal colon tissue (13,14).

Why intervention with calcium is not more effective against colon polyp formation is not fully understood. One thought is that effective chemoprevention requires that there be an adequate level of dietary calcium throughout life. Once early molecular damage has occurred, it is too late to block polyp formation and progression to invasive cancer. Alternatively, calcium supplementation may lack better efficacy because optimal chemoprevention requires additional nutrients along with calcium. Vitamin D, for example, is well-known to be required for calcium uptake and utilization. A vitamin D deficiency is associated with many of the same maladies as a calcium-deficiency including colon cancer (15-18). Other studies have demonstrated that certain trace elements nutritionally associated with calcium may contribute to polyp prevention. For example, the ratio of magnesium to calcium has been shown to be as important as the level of calcium itself for effective colon cancer chemoprevention in mice (19). Other studies have demonstrated anti-tumor activity with copper, manganese and selenium (20,21). Our own past studies have shown that members of the lanthanide family of “rare earth” elements enhance the growth-regulating activity of calcium for colon epithelial cells (22,23). How lanthanide elements affect response to calcium is not known. Certain of the lanthanide elements bind the extracellular calcium-sensing receptor (CaSR), and have a higher affinity than calcium itself (24-27). Since CaSR is a key molecule for transducing epithelial growth-regulatory signals from calcium (28), high-efficiency binding could be expected to affect growth. These, and other potentially important nutrients, would not be provided in a calcium supplement alone.

As part of our effort to understand the role of trace elements in colon polyp chemoprevention, we conducted two long-term (15- and 18-month) studies in mice using Aquamin – a calcium rich, multi-mineral product derived from red marine algae of the *Lithothamnion sp –* as an intervention. Aquamin contains calcium and magnesium in an approximately 12:1 ratio along with detectable levels of 72 additional trace elements (29). In these studies, animals were maintained on a high-fat, low calcium diet with or without the mineral supplement. In the two studies combined (at 12 to 18 months time points), 24 of 70 (34.3%) control mice developed colonic adenomas. Colon polyps were seen in only 2 of 70 (2.9%) mice with Aquamin. Mice on a healthy diet containing a comparable amount of calcium alone (calcium carbonate) developed polyps in 15 of 70 (21.4%) mice (30,31).

In other studies, we examined Aquamin for effects on human colon carcinoma cells in monolayer culture, Aquamin was more effective than calcium alone at suppressing tumor cell growth and inducing differentiation (32,33). Better growth suppression with Aquamin was associated with greater CaSR induction.

Whether the combination of calcium and additional trace elements (be it Aquamin itself or some other mix of appropriate minerals) will, ultimately, prove to be more effective than calcium alone as a colon polyp chemopreventive agent in humans remains to be seen. A problem with translating preclinical findings to results in humans is the low incidence of colon polyp formation and the long lag period between initial molecular changes and outgrowth of observable lesions. Additionally, progression from initial polyp formation to more serious disease is difficult to study experimentally since colonic polyps are removed upon detection. Colonoid culture technology, which is now well-developed (34-37), provides a way to study human colon polyp responses to potentially useful chemopreventive agents under *ex vivo* conditions. In a recent study using human colon adenomas in colonoid culture, we found that supplementation of the culture medium with either calcium alone or Aquamin induced differentiation in the adenomas as indicated by a change in morphology and by increased expression of differentiation-related proteins (38). With Aquamin, features of differentiation were observed at a lower concentration than was required with the equivalent level of calcium alone. With both interventions, differentiation was associated with alterations in several growth-regulating pathways. While these recent studies demonstrate the potential utility of a multi-mineral approach to colon polyp chemoprevention, they do not provide information on specificity – i.e., whether the beneficial activity of Aquamin noted with the adenoma colonoids is unique to the abnormal epithelium or whether changes (perhaps, unwanted) might also be seen in the normal colonic mucosa. As a way to begin addressing that question, we have in the present study compared the effects of Aquamin to calcium alone for ability to modulate growth and differentiation in colonoid cultures derived from specimens of histologically normal human colonic mucosa.

## MATERIALS AND METHODS

### Aquamin

Aquamin is a calcium-rich, magnesium-rich multi-mineral product obtained from the skeletal remains of the red marine algae, *Lithothamnion sp* (29) (Marigot Ltd, Cork, Ireland). Aquamin contains calcium and magnesium in a ratio of approximately (12:1), along with measurable levels of 72 other trace minerals (essentially all of the minerals algae fronds accumulate from the deep ocean water). Mineral composition was established via an independent laboratory (Advanced Laboratories; Salt Lake City, Utah) using Inductively Coupled Plasma Optical Emission Spectrometry (*ICP*-*OES*). Supplement Table 1 from our recently published study with adenoma colonoids provides a complete list of elements detected in Aquamin and their relative amounts (38). Aquamin is sold as a dietary supplement (GRAS 000028) and is used in various products for human consumption in Europe, Asia, Australia, and North America. A single batch of Aquamin®-Soluble was used for this study. Calcium Chloride 0.5 M solution (PromoCell GmbH, Heidelberg, Germany) was used as a source of calcium.

### Tissue samples for colonoid culture

Histologically normal colon tissue (2-mm biopsies) were provided by five subjects using unpreped flexible sigmoidoscopy. The five subjects were from a group defined as at “increased risk for colon cancer based on a personal history of colonic polyps or a family (first degree blood relative) history of colon cancer.” The study was approved by the Institutional Review board (IRBMED) at the University of Michigan (HUM 00076276), and all subjects provided written informed consent prior to biopsy. These subjects were also screened for use of calcium or vitamin D supplements, and any supplement use was stopped two weeks prior to the biopsy. Supplement Table 1 provides demographics information.

### Establishment of colonoids from normal human colonic mucosa

Colonoid cultures were established from histologically-normal colon tissue based on our recently described methods (37,39). Briefly, biopsies were finely minced on ice using a #21 scalpel and seeded into matrigel. During the expansion phase, colonoids were incubated in L-WRN medium (10% fetal bovine serum), which provides a source of Wnt3a, R-spondin-3 and Noggin. The medium was supplemented with small molecule inhibitors: 500nM A 83-01 (Tocris), a TGF-β inhibitor, 10µM SB 202190 (Sigma), a p38 inhibitor and 10 µM Y27632 as a ROCK inhibitor, (Tocris) (40). For the first 10 days of culture, the medium was also supplemented with 2.5µM CHIR99021 (Tocris). Established colonoids were interrogated in a mix of L-WRN culture medium diluted 1:4 with KGM Gold, a serum-free, calcium-free culture medium (Lonza). The final serum concentration was 2.5% and calcium concentration was 0.25 mM. This “control treatment medium” was compared to the same medium supplemented with calcium to a final concentration of 1.5 – 3.0 mM. Calcium was provided alone as calcium chloride or as Aquamin formulated to contain the equivalent amount of calcium.

### Phase-contrast microscopy

Colonoids were evaluated by phase-contrast microscopy (Hoffman Modulation Contrast - Olympus IX70 with a DP71 digital camera) for change in size and shape during the in-culture part of the study.

### Histology, immunohistology and Morphometric analysis

At the end of the in-life phase, colonoids were isolated from Matrigel using 2mM EDTA and fixed in 10% formalin for 1 hour. Fixed colonoids were suspended in HistoGel (Thermo Scientific) and then processed for histology (i.e., hematoxylin and eosin staining) or for immunohistology. For this, freshly-cut sections (5-6 microns) were rehydrated, and subjected to heat-induced epitope retrieval with high pH or low pH FLEX TRS Retrieval buffer (Agilent Technologies, 154 #K8004; Santa Clara, CA) for 20 minutes. After peroxidase blocking, antibodies were applied at appropriate dilutions at room temperature for 30 or 60 minutes (Supplement Table 2). The FLEX HRP EnVision System (Agilent Technologies) was used for detection with a 10-minute DAB chromagen application. Supplement Table 2 provides a list of antibodies used and their source.

The sections of immunostained colonoid tissue on glass slides were digitized using the Aperio AT2 whole slide scanner (Leica Biosystems) at with a resolution of 0.5µm per pixel with 20x objective. These scanned images were housed on a server and accessed using Leica Aperio eSlide Manager (Version 12.3.2.5030), a digital pathology management software. The digitized histological sections were viewed and analyzed using Aperio ImageScope (Version 12.3.3.5048), a slide viewing software. Brightfield Immunohistochemistry Image Analysis tools (Leica) were used to quantify different immunostains used in this study. Aperio Nuclear Algorithm (v9) was used for proliferation markers (Ki67) quantification. This algorithm measures intensity of the nuclear staining and separate those into very intense to no nuclear staining (3+, 2+, 1+ and 0 respectively). Nuclei with 3+ intensity were used here for comparison. The Aperio Positive Pixel Count Algorithm (v9) was used to quantify differentiation marker expression. It quantifies number and the intensity of pixels of a specific stain in a digitized image. Positivity was calculated with respective numbers of strong positive and positive pixels against total pixels.

### Scanning electron microscopy (SEM) and transmission electron microscopy (TEM)

Colonoid specimens were fixed *in situ* in 2.5 percent glutaraldehyde in 0.1 M Sorensen’s buffer, pH 7.4, overnight at 4°C. After subsequent processing for SEM or TEM as described previously (41), samples for SEM were then mounted on stubs, allowed to off-gas in a vacuum desiccator for at least two hours and sputter coated with gold. Samples were examined with an Amray 1910 FE Scanning Electron Microscope and digitally imaged using Semicaps 2000 software. For TEM, ultra-thin sections were examined using a Philips CM100 electron microscope at 60 kV. Images were recorded digitally using a Hamamatsu ORCA-HR digital camera system operated with AMT software (Advanced Microscopy Techniques Corp., Danvers, MA).

### Differential proteomic analysis

Colonoids were isolated from Matrigel using 2mM EDTA for 15 minutes and then exposed to Radioimmunoprecipitation assay (RIPA)-lysis and extraction buffer (Thermo Scientific, Rockford, IL) for protein isolation. Proteomic experiments were carried out in the Proteomics Resource Facility (PRF) in the Department of Pathology at the University of Michigan, employing mass spectrometry-based Tandem Mass Tag (TMT, ThermoFisher Scientific). Fifty micrograms of colonoid protein from each condition was digested with trypsin and individually labeled with one of the 10 isobaric mass tags following the manufacturer’s protocol. After labeling, equal amounts of peptide from each condition were mixed together. In order to achieve in-depth characterization of the proteome, the labeled peptides were fractionated using 2D-LC (basic pH reverse-phase separation followed by acidic pH reverse phase) and analyzed on a high-resolution, tribrid mass spectrometer (Orbitrap Fusion Tribrid, ThermoFisher Scientific) using conditions optimized at the PRF. MultiNotch MS3 approach (42) was employed to obtain accurate quantitation of the identified proteins/peptides. Data analysis was performed using Proteome Discoverer (v 2.1, ThermoFisher). MS2 spectra were searched against SwissProt human protein database (release 2016-11-30; 42054 sequences) using the following search parameters: MS1 and MS2 tolerance were set to 10 ppm and 0.6 Da, respectively; carbamidomethylation of cysteines (57.02146 Da) and TMT labeling of lysine and N-termini of peptides (229.16293 Da) were considered static modifications; oxidation of methionine (15.9949 Da) and deamidation of asparagine and glutamine (0.98401 Da) were considered variable. Identified proteins and peptides were filtered to retain only those that passed ≤2% false-discovery rate (FDR) threshold of detection. Quantitation was performed using high-quality MS3 spectra (Average signal-to-noise ratio of 9 and <40% isolation interference). Differential protein expression between conditions, normalizing to control (0.25mM calcium) for each subject’s specimens separately was established using edgeR (43). Then, results for individual proteins from the three subjects were averaged. Proteins names were retrieved using Uniprot.org, and reactome V66 (reactome.org) was used for pathway enrichment analyses (44). Only Proteins with a ≤2% FDR confidence of detection were included in the analyses. The initial analysis was targeted towards differentiation, barrier-related and cell-matrix adhesion proteins. Follow-up analysis involved an unbiased proteome-wide screen of all other proteins modified by Aquamin or by calcium alone in relation to control. String database (string-db.org) was used to identify interactions between differentially-expressed proteins.

### Statistical analysis

Means and standard deviations were obtained for discrete morphological and immunohistochemical features as well as for individual proteins. Groups were analyzed by ANOVA followed by student t-test (two-tailed) for unpaired data. Pathways enrichment data reflect Reactome-generated p-values based on the number of entities identified in a given pathway as compared to total proteins responsible for that pathway. Data were considered significant at p<0.05.

## RESULTS

### Effects of calcium alone and Aquamin on structure of colonoids derived from histologically normal tissue

Histologically normal colon tissue was established in colonoid culture and incubated for a two-week period in control medium (0.25 mM calcium) or in the same culture medium supplemented with calcium to a final concentration of 1.5 – 3.0 mM or with Aquamin to provide the same levels of calcium. At the end of the incubation period, colonoids were examined for gross appearance by phase-contrast and scanning electron microscopy and for microscopic appearance after sectioning and staining with hematoxylin and eosin.

Representative phase-contrast images from day-14 cultures are shown in Figure 1a-c. Colonoids maintained under control conditions (0.25 mM calcium) appeared as either thin-walled, translucent, cystic structures (arrows) that were mostly spherical or thick-walled structures with a variety of shapes (Figure 1a). When colonoids were maintained in the same medium but supplemented with calcium to a final concentration of 1.5 mM (provided either as calcium chloride or as Aquamin) (Figure 1b and 1c), there was no significant alteration in morphology. Both thin-walled, spherical structures and thick-walled structures were apparent. Additional colonoids (not shown) were treated with calcium (alone or as Aquamin) at 2.1 and 3.0 mM. Colonoids maintained at these higher calcium levels were similar in appearance to those treated with 1.5 mM calcium. Cultures from all five subjects’ tissue were similar in appearance.

**Figure 1.**
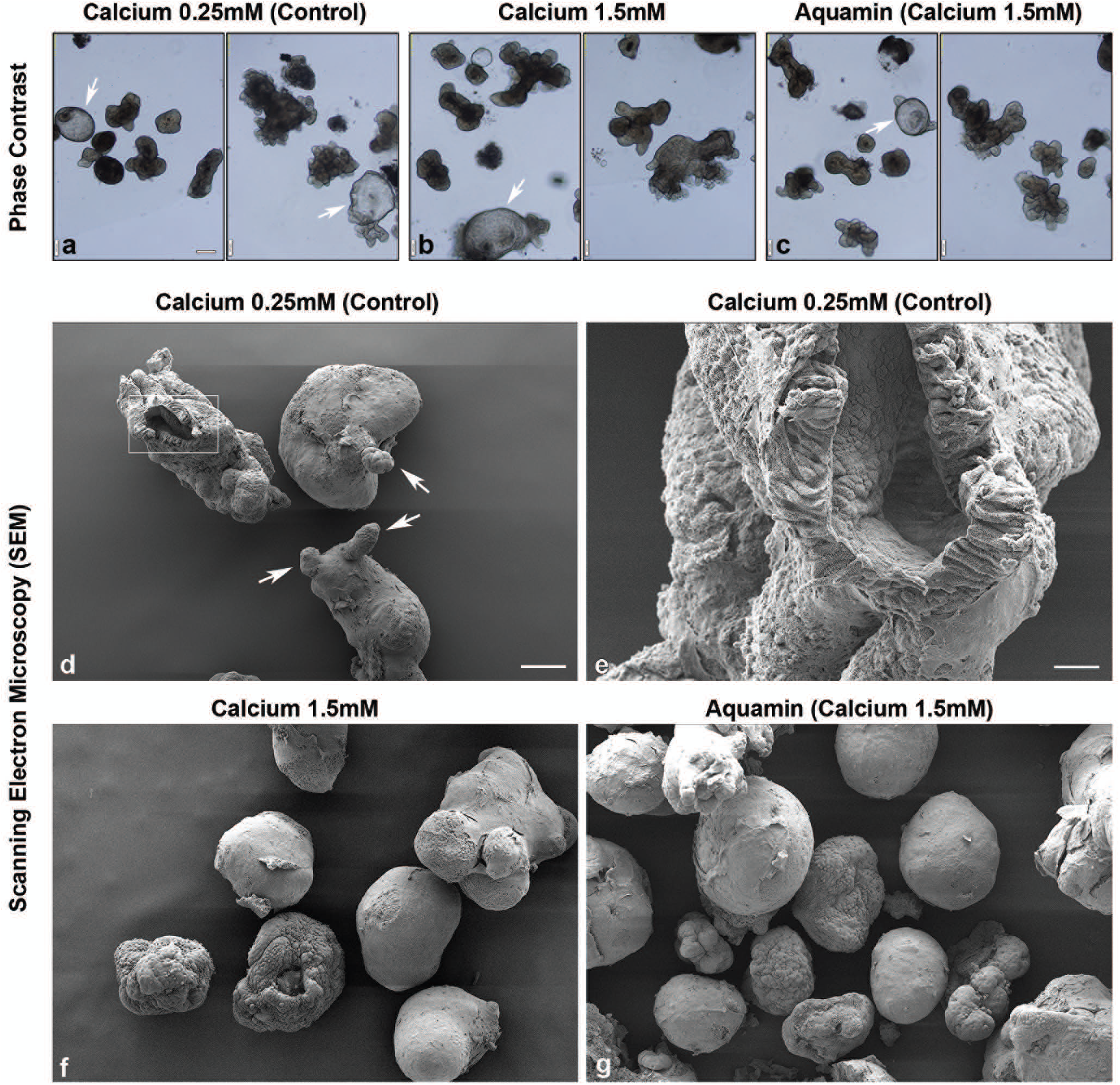
Colonoid appearance in culture: Phase contrast and scanning electron microscopy. At the end of the incubation period, intact colonoids were examined. Under phase contrast microscopy **(a-c)**, colonoids were present either as thin-walled, translucent cystic structures (arrows) or as thick-walled structures with few surface buds. They were similar in appearance under all conditions. Scanning electron microscopy confirmed the presence of smooth surface and few buds (arrows) in colonoids maintained under low-calcium conditions **(d)**. At higher magnification **(e),** colonoids were shown to consist of hollow structures with a central lumen surrounded by columnar epithelial cells. Under high-calcium conditions **(f and g)**, colonoids were similar to those maintained in the low-calcium medium but without the surface buds. Bar for a, b and c =200μm; bar for d, f and g =100μm; bar for e = 20μm.

Scanning electron microscopy provided more precise evaluation of colonoid morphology (Figure 1d-g). Under control (low-calcium) conditions (Figure 1d), colonoids had a spherical or oblong shape with a smooth surface. They were typically 100-200 μm in diameter. Most individual colonoids appeared to be “wrapped” with the matrix support, but in some cases, there was no visible matrix and the cells, themselves, formed the surface of the structure. Occasional small projections (buds) protruded from the colonoid surface (white arrows). In places, the colonoid surface was open (insert), allowing for visualization of the interior of the structure. At higher magnification (Figure 1e), it could be seen that the colonoid was hollow, with a central lumen surrounded by columnar epithelial cells (arrows). Colonoids maintained in the presence of 1.5 mM calcium (either alone or as Aquamin) (Figure 1f and 1g) were similar in appearance to the colonoids maintained in low-calcium medium. The one difference was that the small projections (buds) present in low-calcium colonoids were not seen at the higher calcium levels.

Figure 2a-c demonstrates microscopic appearance of colonoids maintained under the same three conditions. Under all three conditions, colonoids were present as a mix of thin-walled crypts with spherical lumens, or as thick-walled crypts with lumens of varying shapes and sizes. The epithelial cells in the thin-walled crypts were mostly cuboidal in shape while the cells in the thick-walled structures were a mix of cells with cuboidal or columnar morphology. Although there were no dramatic differences between the treatment groups and control in microscopic appearance, when lumen diameter and wall-thickness were quantified from hematoxylin and eosin stained sections, an increase in both parameters was observed under high-calcium conditions as compared to control (Figure 2d and 2e). Differences between colonoids exposed to calcium alone versus Aquamin were minimal.

**Figure 2.**
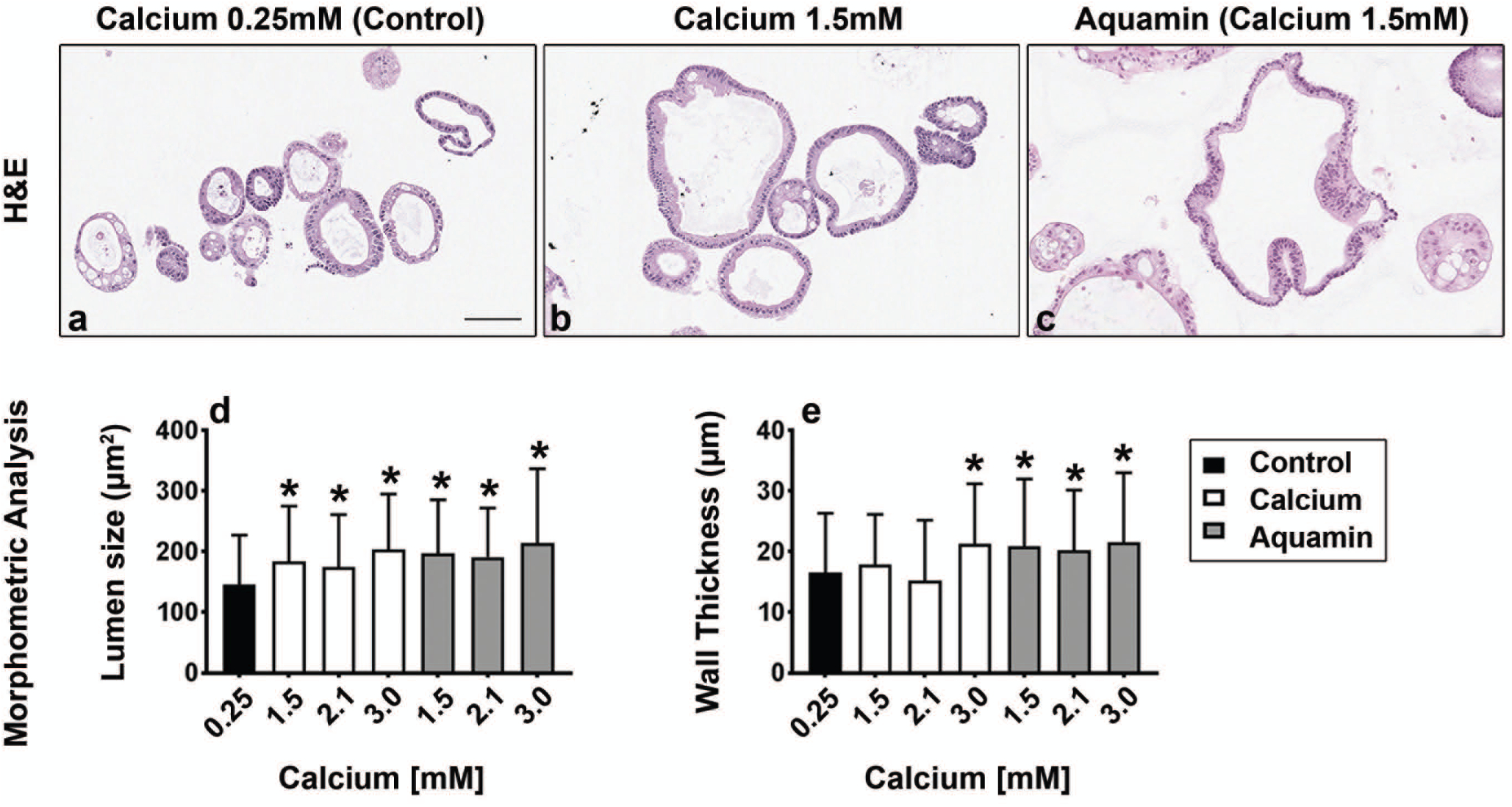
Colonoid appearance in culture: Microscopic features. At the end of the incubation period, colonoids were examined by light microscopy after staining with hematoxylin and eosin **(a-c)**. Under all three conditions, the colonoids were found to be crypts of varying size with a single layer of epithelial cells surrounding a central lumen. Under control conditions **(a)**, small crypts (with as few as 20 cells in cross section) were seen. In the presence of 1.5 mM calcium alone **(b)** or with Aquamin providing 1.5 mM calcium **(c)**, larger crypts made up of columnar epithelial cells surrounding a large, often irregular-in-shape lumen were seen. Goblet cells were apparent. Lumen size and wall thickness **(d and e)** are shown in the accompanying bar graphs. Means and standard deviations are based on 74-123 individual crypts per condition. Asterisks indicate statistical significance from control at p<0.05 level. Bar=100μm.

### Proliferation and differentiation marker expression patterns

Quantitative immunohistology was used to assess Ki67 expression as a proliferation marker and CK20 as a marker of epithelial cell differentiation (Figure 3). Ki67 expression was decreased modestly with intervention but it was significant with higher concentration of either intervention (Figure 3a-c). CK20 was strongly positive in virtually all of the colonoids regardless of treatment (Figure 3d-f).

**Figure 3.**
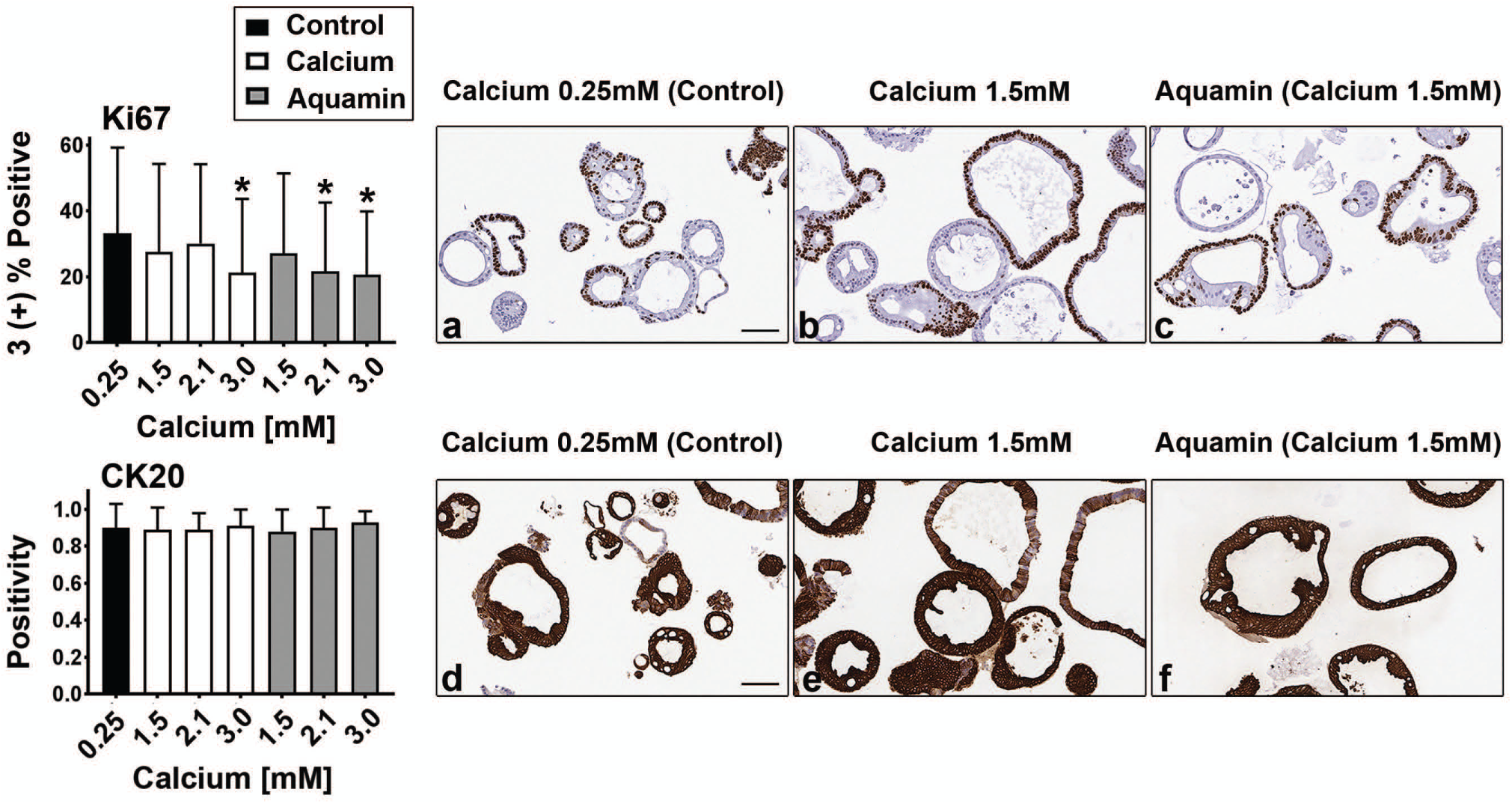
Proliferation and differentiation marker expression: Immunohistology. At the end of the incubation period, colonoids were examined after immunostaining. **Upper panels (a, b and c)**: Ki67 expression. A mix of positive and negative staining was observed under all conditions. **Lower panels (d, e and f)**: CK20 expression. Most crypts, regardless of size or shape were strongly positive. Morphometric analysis is shown in the accompanying bar graphs. Ki67 values (based on nuclear algorithm v9) are means and standard deviations based on 58-114 individual crypts per condition. Asterisks indicate statistical significance from control at p<0.05 level. CK20 values represent positivity (measured using Positive Pixel Value v9). Means and standard deviations based on 78-126 individual crypts per condition. Asterisks indicate statistical significance from control at p<0.05 level. Bars=100μm

### Proteomic analysis: Up-regulation of proteins involved in differentiation and related functions

We utilized a mass spectrometry-based approach to identify differentiation-related proteins expressed in colonoid cultures and assess the effects of calcium alone and Aquamin on expression levels relative to control. Table 1A is a compilation of known differentiation-related proteins as well as proteins involved in cell-cell adhesion, cell-matrix adhesion and barrier function. Average fold-change values across the three specimens with each intervention at day-14 are compared to control conditions. As can be seen from the table, there was an increase in CK20 expression along with an increased level of epiplakin – a protein required for intermediate filament network formation and differentiation (45). A number of other keratins (as well as proteins such as hornerin and filaggrin) were also detected in the cell lysates (not shown). Some of these showed a modest increase with intervention in one or two subjects, but most did not reach statistical significance because they were not up-regulated in colonoids from all three subjects.

**Table 1A.**
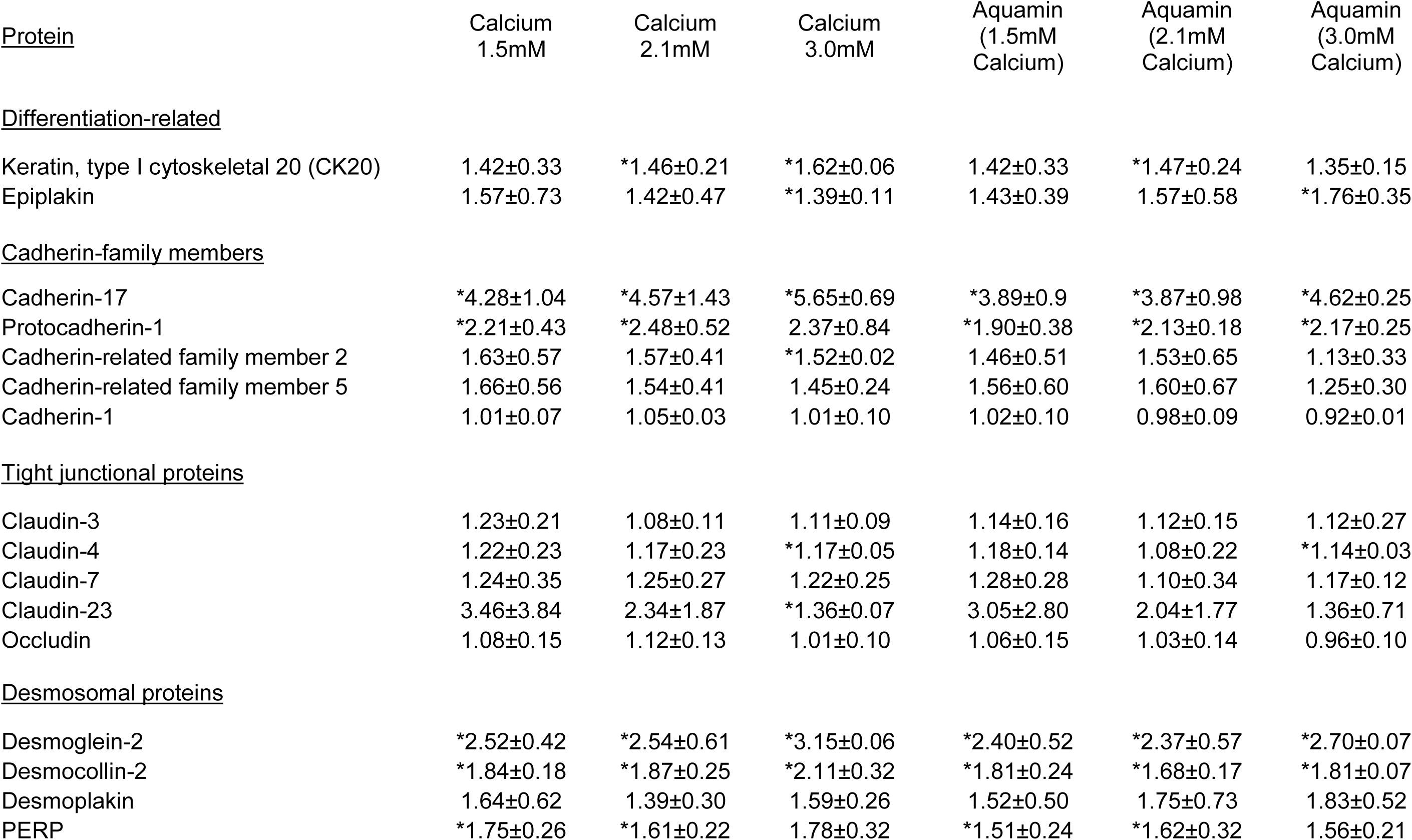
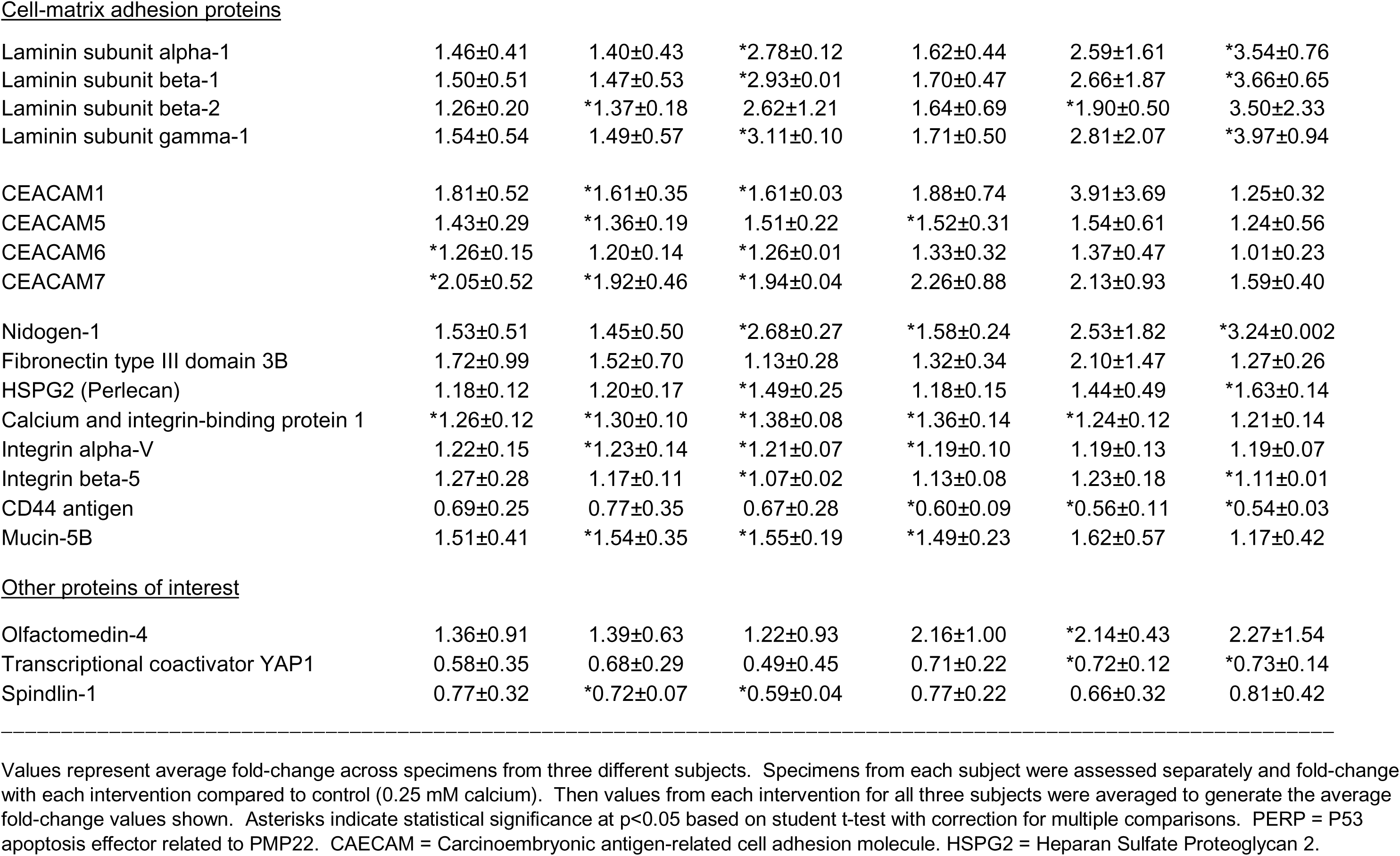
Differentiation-related proteins.

The same lysates were examined for proteins that make up *adherens* junctions (cadherin family members), tight junctions (claudins and occludin) and desmosomes (desmoglein-2, desmocollin-2, desmoplakin and p53 apoptosis effector related to PMP-22 [PERP]). Several cadherin family members were strongly and consistently up-regulated as were desmosomal proteins (Table 1A). In contrast, tight junction protein up-regulation was more modest. Only claudin-23 was substantially increased, and this was due, primarily, to up-regulation in one specimen only (Table 1A).

Table 1A also demonstrates that several proteins involved in cell-matrix adhesion were substantially elevated in samples treated with calcium alone or Aquamin as compared to control. Among these were laminin subunits, several carcinoembryonic antigen-related cell adhesion molecules (CAECAMs), nidogen, fibronectin type III domain protein and perlecan (basement membrane specific heparin sulfate proteoglycan-2). Integrin alpha V and integrin beta 5 subunits were also up-regulated with the two interventions (Table 1A) while virtually all of the other alpha and beta integrin subunits showed little difference with calcium alone or with Aquamin treatment (not shown). In contrast to these up-regulated proteins, CD44, a hyaluronan receptor associated with multiple cell functions including cell motility was down-regulated with both interventions, but more strongly with Aquamin (Table 1A). High CD44 expression has been linked to cancer metastasis (46).

Known interactions among the proteins listed in Table 1A are presented using String database (Supplement Figure 1). As seen, the cell-cell and cell-matrix adhesion proteins cluster together. These clusters interact with each other through cadherins (primarily cadherin 17). Table 1B highlights the top 24 pathways associated with the proteins presented in Table 1A. Not surprisingly, most of these pathways involve the extracellular matrix, cell-cell junctional organization and cell-cell communication.

**Table 1B.** Top pathways involved with proteins presented in table 1A.

| **<u>Pathway name</u>** | **<u>Entities</u>**<br>**<u>pValue</u>** | **<u>Mapped entities</u>** |
| --- | --- | --- |
| Laminin interactions | $6.7 \times 10^{-12}$ | LAMB1;ITGAV;LAMB2;LAMA1;HSPG2;LAMC1;NID1 |
| Extracellular matrix organization | $7.4 \times 10^{-11}$ | LAMB1;CEACAM6 ;ITGAV;LAMB2;LAMA1;ITGB5;CDH1;CEACAM1;HSPG2;LAMC1;NID1; CD44 |
| Non-integrin membrane-ECM interactions | $7.1 \times 10^{-10}$ | LAMB1;ITGAV;LAMB2;LAMA1;ITGB5;HSPG2;LAMC1 |
| ECM proteoglycans | $4.0 \times 10^{-9}$ | LAMB1;ITGAV;LAMB2;LAMA1;ITGB5;HSPG2;LAMC1 |
| Apoptotic cleavage of cell adhesion proteins | $4.9 \times 10^{-8}$ | DSG2;OCLN;DSP;CDH1 |
| Cell-cell junction organization | $5.7 \times 10^{-8}$ | CLDN23;CLDN3 ;CLDN4 ;CDH17;CLDN7 ;CDH1 |
| Cell junction organization | $4.3 \times 10^{-7}$ | CLDN23;CLDN3 ;CLDN4 ;CDH17;CLDN7 ;CDH1 |
| Tight junction interactions | $2.6 \times 10^{-6}$ | CLDN23;CLDN3 ;CLDN4 ;CLDN7 |
| Cell-Cell communication | $3.1 \times 10^{-6}$ | CLDN23;CLDN3 ;CLDN4 ;CDH17;CLDN7 ;CDH1 |
| Apoptotic cleavage of cellular proteins | $6.6 \times 10^{-6}$ | DSG2;OCLN;DSP;CDH1 |
| Integrin cell surface interactions | $7.1 \times 10^{-6}$ | ITGAV;ITGB5;CDH1;HSPG2; CD44 |
| Apoptotic execution phase | $2.3 \times 10^{-5}$ | DSG2;OCLN;DSP;CDH1 |
| Formation of the cornified envelope | $5.2 \times 10^{-5}$ | DSG2;DSC2;DSP;KRT20;PERP |
| Fibronectin matrix formation | $1.7 \times 10^{-4}$ | CEACAM6 ;CEACAM1 |
| Keratinization | $5.7 \times 10^{-4}$ | DSG2;DSC2;DSP;KRT20;PERP |
| L1CAM interactions | $5.7 \times 10^{-4}$ | LAMB1;ITGAV;LAMA1;LAMC1 |
| Apoptosis | 0.002 | DSG2;OCLN;DSP;CDH1 |
| Programmed Cell Death | 0.002 | DSG2;OCLN;DSP;CDH1 |
| Syndecan interactions | 0.003 | ITGAV;ITGB5 |
| Adherens junctions interactions | 0.005 | CDH17;CDH1 |
| Developmental Biology | 0.005 | LAMB1;DSG2;ITGAV;DSC2;LAMA1;DSP;KRT20;PERP;LAMC1 |
| RUNX1 regulates expression of components of tight junctions | 0.02 | OCLN |
| PECAM1 interactions | 0.04 | ITGAV |
| TP53 regulates transcription of cell death genes | 0.04 | PERP |

Reactome database (v66) and Uniprot are the source of information.

Although our primary focus was on proteins involved in differentiation and related functions, we also used the proteomic approach to conduct a non-biased search of all proteins up-regulated or down-regulated by either intervention. Average increase or decrease values across the three specimens (representing proteins altered by 1.8-fold or greater with <2% FDR in at least one specimen) are presented in Figure 4. The Venn plots shown in the left-portion of the figure demonstrate a substantial concurrence between proteins up- or down-regulated by the two interventions at comparable calcium levels and the scatter plots in the center show the relationship between the degree of up- or down-regulation in the overlapping proteins. The Venn plots shown in the right-hand portion of the figure demonstrate the distribution of proteins among the three specimens. Interestingly, only one protein – cadherin-17 – met the criterion of being increased by greater than 1.8-fold with both interventions at all concentrations in all three specimens.

**Figure 4.**
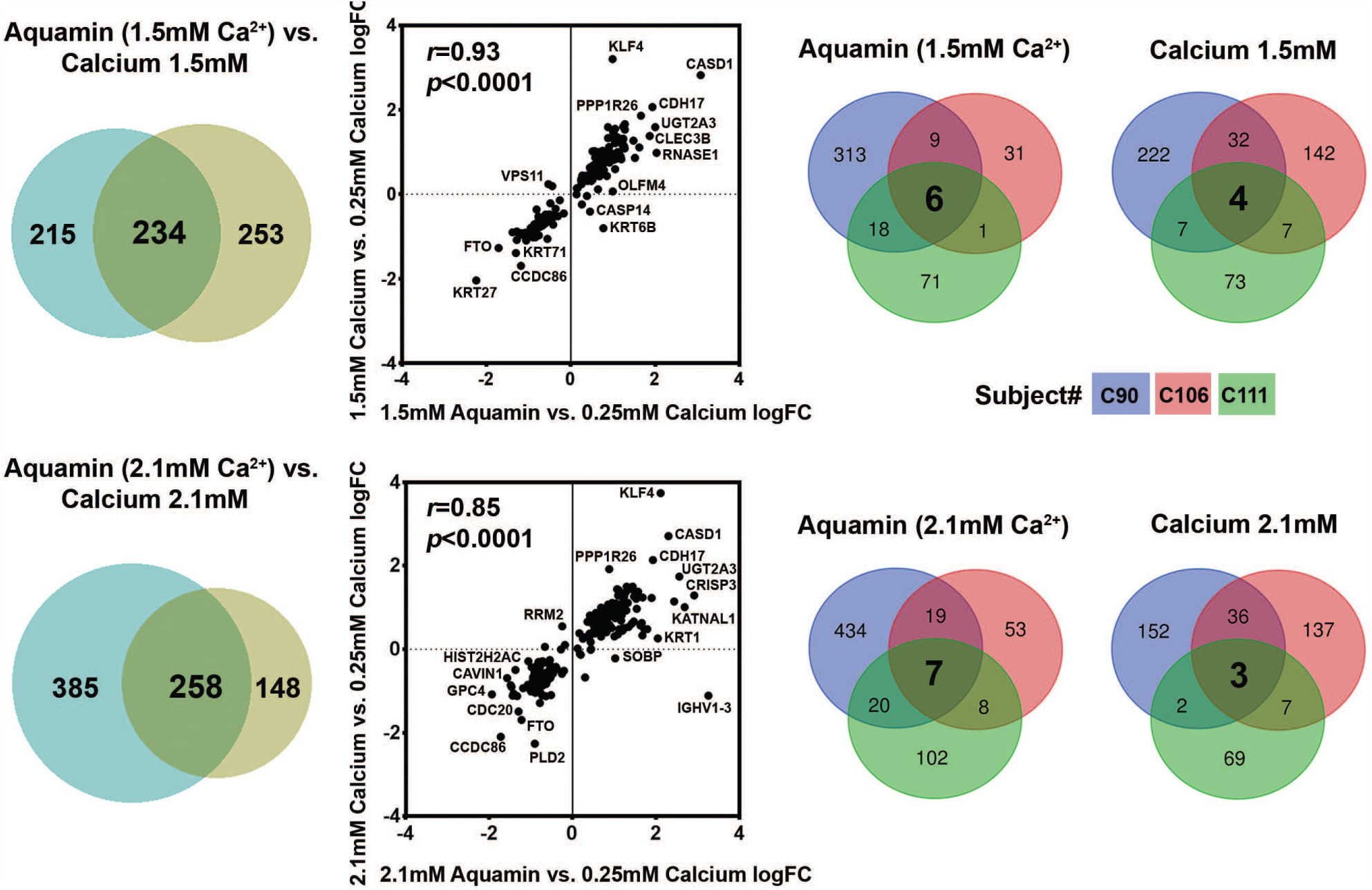
Proteomic analysis. At the end of the incubation period, lysates were prepared for proteomic analysis. **Left**: Venn plots showing proteins altered (increased or decreased) by an average of 1.8-fold or greater across the three specimens in response to calcium alone or to Aquamin with a comparable amount of calcium, and the overlap between pairs of interventions. **Center**: The scatterplots demonstrate quantitative relationships between individual proteins altered by each pair of interventions. Correlation between the two interventions at both concentrations were significant (p<0.0001). **Right**: Venn plots showing proteins altered (increased or decreased) by an average of 1.8-fold or greater with each intervention and the overlap among individual specimens. This is to show the variability in subjects and a response to the intervention.

Several individual proteins of interest (in addition to the ones discussed above) were identified with an unbiased approach. These are presented in Supplement Tables 3A and 4A). Among up-regulated proteins was vitamin D-binding protein, the major transporter of vitamin D in the circulation. This protein is responsible for carrying vitamin D into cells (47) and is, therefore, critical to calcium uptake and utilization at the cellular level. Intestinal alkaline phosphatase and 15-hydroxyprostaglandin dehydrogenase were two additional proteins of interest that were substantially up-regulated. Elevated expression of these proteins are associated with inflammation-suppression in the gastrointestinal tract (48-51). They provide potential links between improved barrier function and decreased inflammation. Among down-regulated moieties were proteins related to growth - i.e., proliferating cell nuclear antigen (PCNA) (52) and nucleophosmin (NPM) (53,54). Supplement Tables 3B and 4B presents pathways associated with the up-regulated and down-regulated proteins.

Finally, we searched the list of up-regulated and down-regulated proteins for other moieties of interest and for proteins that had been shown in our previous study (38) to be strongly affected by calcium alone or Aquamin in colonoid cultures established from large, premalignant adenomas. Among the proteins of interest identified in adenomas were moieties associated with growth-regulating pathways; i.e. NF2/merlin, BRCA-related protein, members of the histone 1 family and olfactomedin-4. Among down-regulated proteins were metallothionine 1E and 1H. Of these, only olfactomedin-4 was sufficiently abundant to reach threshold level in normal colonoids. It was modestly up-regulated by calcium alone, but more highly induced by Aquamin (1.22 - 1.39-fold increase with calcium - not significant; and 2.14 – 2.27-fold with Aquamin; p<0.05) (Table 1A). Olfactomedin-4 is of interest because while it is clearly responsive to differentiation-inducing interventions, it is also expressed in cells in concurrence with other stem cell markers (55). Also of interest, two transcription enhancers associated with dysregulated growth (YAP-1 and spindlin-1) (56,57) were both down-regulated with calcium and Aquamin (Table 1A). YAP-1 is a down-stream target in the HIPPO signaling pathway. NF2, a potent inhibitor of HIPPO signaling, was strongly up-regulated by Aquamin in adenoma colonoids (38).

### Immunohistological and ultrastructural assessment of barrier proteins

Based on proteomic findings, we chose three proteins – cadherin-17, claudin-23 and desmoglein-2 for immunohistological assessment. All three proteins were expressed under control conditions; a mixture of cytoplasmic and cell surface staining was seen (Figure 5). With cadherin-17 and desmoglein-2, a clear shift from cytoplasmic to cell surface could be seen with either intervention (calcium at 1.5 mM or Aquamin with equivalent calcium). Alterations in protein distribution was less evident with claudin-23.

**Figure 5.**
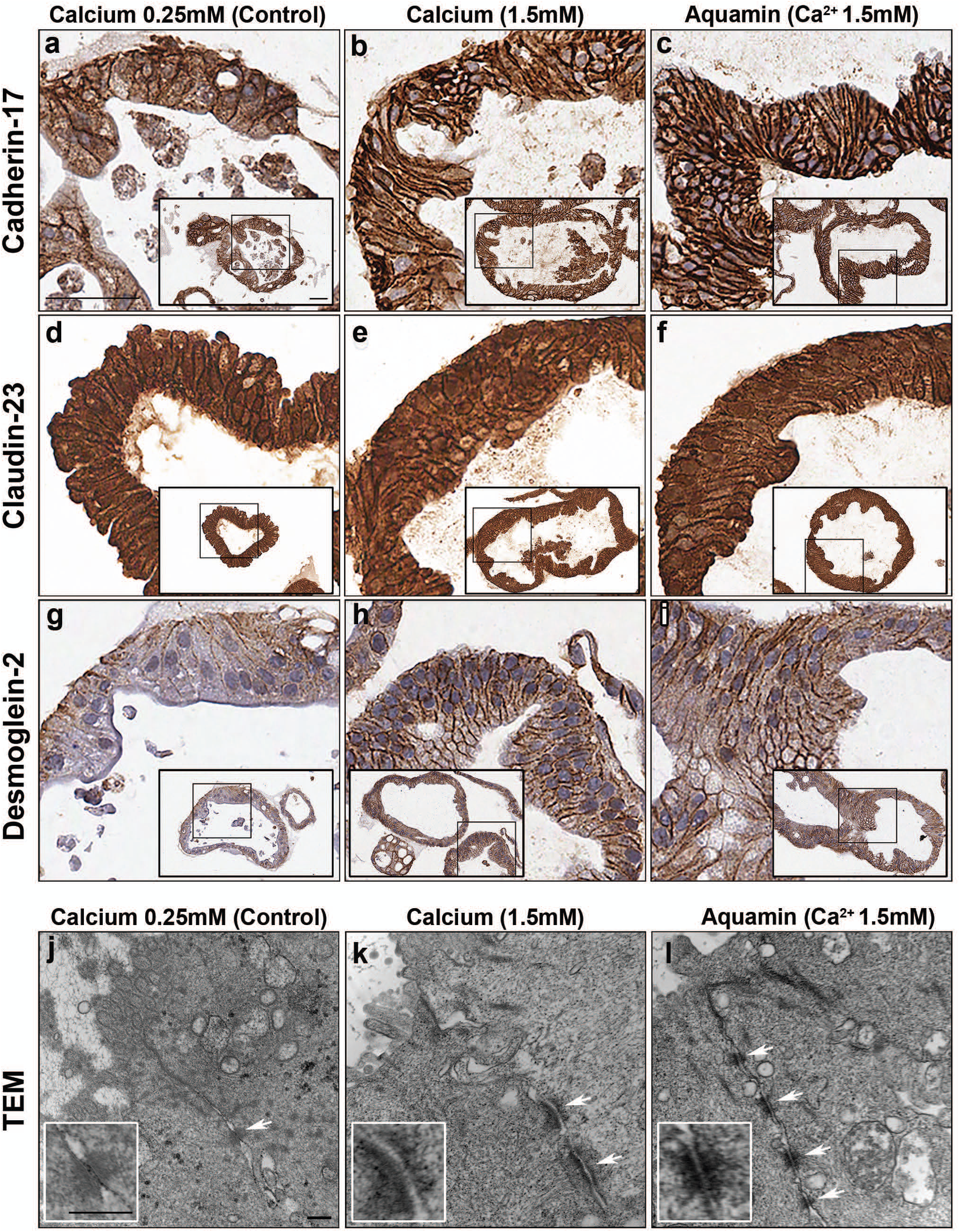
Cell surface components: At the end of the incubation period, colonoids were examined after immunostaining (**upper panels**) and transmission electron microscopy (**lower panels**). Cadherin-17 **(a, b and c)**; claudin-23 **(d, e and f)**; desmoglein-2 **(g, h and i)**: All three proteins are visible in colonoids under all conditions by immunohistology. With cadherin-17 and desmoglein-2, there is a clear shift from cytoplasmic to surface with intervention. With claudin-23, high surface staining is apparent under all conditions. Bars for (a-i) main panel and inset=50μm. Lower panels: transmission electron microscopy. A higher density of desmosomes along the lateral surface can be seen with intervention (at 10,000X). Bar=100 μm (Insets bar=200 μm).

In a final set of studies, colonoid specimens from four subjects were examined by transmission electron microscopy. The focus was on regions of cell-cell contact. Electron microscopy demonstrated differences in desmosome expression in response to intervention as compared to control (Figure 5). Under low-calcium conditions, desmosomes were few in number, size and electron density. We rarely saw more than a single desmosome in any high-power (10,000X) image. In contrast, colonoids maintained in medium containing 1.5 mM calcium (alone or as Aquamin) demonstrated large numbers of desmosomes between cells. Individual desmosomes were wider and more electron dense. Well-organized intermediate filaments connected to the desmosomes were apparent in places. Colonoids maintained in calcium alone and colonoids treated with Aquamin were indistinguishable from each other. Findings were similar with all subjects.

Tight junctional complexes were also evident by electron microscopy. These were present at the luminal surface between virtually every cells, regardless of whether the colonoids were cultured under control conditions or exposed to either intervention (not shown). This is in contrast to what was observed previously in adenoma colonoids. In the adenoma colonoids, there were few detectable tight junctions under low-calcium conditions. However, when exposed to higher calcium concentrations, tight junctions were apparent (38).

## DISCUSSION

In a recent study, we characterized the response of human colonic adenomas in colonoid culture to calcium supplementation (38). When tumor colonoids were maintained under low calcium (0.15 mM) conditions, they were highly-proliferative. Numerous small buds continued to form on the surface as long as the colonoids were in culture. When the structures were fragmented at each subculture, new colonoids formed from the existing buds with high efficiency. At the histological level, the buds were found to be spherical crypts comprised of cells with cuboidal morphology surrounding a small circular central lumen. Most of the cells (>80%) were Ki67-positive and there was little CK20 expression. When the same colonoids were exposed to medium containing 1.5 mM calcium (or higher), fewer new buds formed, and existing buds grew larger in size. At the histological level, the large buds had the appearance of differentiated crypts – i.e., with a large, irregular lumen surrounded by columnar epithelial cells, some with visible goblets. Accompanying the morphological alterations were a reduction in Ki67 (proliferation marker) expression and an increase in expression of CK20 (differentiation marker).

In the present study, colonoid cultures from histologically-normal colon tissue were examined under low-calcium (0.25 mM) conditions and compared to colonoids treated with calcium concentrations (1.5 - 3.0 mM) that induced differentiation and suppressed proliferation in adenoma colonoids. Compared to what was observed with adenoma colonoids, the normal tissue-derived structures had several noticeable differences. Most importantly, while normal colonoids did, indeed, demonstrate increased differentiation in response to supplementation with calcium or Aquamin, they also showed histological features of differentiation and differentiation-related surface marker expression profiles when maintained in the low-calcium environment. What accounts for the higher sensitivity of the normal colonoids to low ambient calcium is unclear. Previously it has been shown that CaSR, a critical regulator of calcium responses in epithelial cells (28) is highly-expressed in the normal colonic mucosa but reduced or missing completely in colon tumors (24,25). Differences in CaSR expression levels could account for the findings described here. Regardless of mechanism, the potential of epithelial cells in the adenomas to undergo differentiation in response to higher calcium concentrations suggests that even after the premalignant tumors have reached the “large adenoma” size, there is still potential for calcium-supplementation to have a pro-differentiating effect. This is of interest because it has been shown in past studies with both human colon adenomas and human colon carcinomas that a histologically-differentiated presentation is a favorable prognostic factor (58-62).

A long term goal of our work has been to determine if the inclusion of additional trace elements along with calcium can improve on the efficacy of calcium alone as a colon polyp chemopreventive agent. Utilizing a natural product, i.e., - Aquamin - that contains a high concentration of magnesium as well as measurable levels of 72 additional trace elements in addition to calcium, we have shown improved colon polyp prevention in long-term animal studies (30,31) compared to calcium alone. Better epithelial growth suppression in cell culture (32,33) has also been observed. Consistent with these findings, Aquamin (at a level providing only 0.15 mM calcium) induced features of differentiation in adenoma colonoids not seen with calcium alone (38). Aquamin was also more effective than calcium alone in up-regulating proteins with growth-suppressing function. Among these were NF2, a cancer suppressor protein in the P21 pathway and the hippo signaling pathway (63-66) and various H1 histone family members. Aquamin was also more effective in down-regulating metallothionines than was calcium alone. Here we show in normal tissue colonoids greater induction of olfactomedin-4 with Aquamin and greater down-regulation of CD44 with the multi-mineral product than with calcium alone. Most of the other barrier proteins were comparably affected. The overall lack of widespread differences between the two interventions in the normal colonic cultures makes it unlikely that Aquamin will have a negative impact on the normal colonic mucosa when used as a chemopreventive agent.

Although intervention with either calcium alone or Aquamin produced only modest change in the gross and histological appearance of normal tissue-derived colonoids, there was a strong induction of several proteins that contribute to cellular adhesive functions, barrier formation and tissue integrity. Among these were laminin subunits, nidogen, a fibronectin domain protein, Integrin alpha V and beta 5 subunits and a number of CAECAMS (cell-matrix adhesion), cadherin family members (*adherens* junctions), claudins (tight junctions) and desmosomal proteins including desmoglein-2, desmocollin-2, desmoplakin and PERP. At the ultrastructural level, a large numbers of actual desmosomes along the lateral cell-cell boundary could be seen in colonoids exposed to the interventions. These structures were sparse in colonoids cultured under low-calcium conditions.

The implication of these findings goes beyond the normal colonocyte – tumor cell transformation process. All of these cell-cell and cell-matrix adhesion proteins are needed for barrier formation and tissue integrity. Tight junctions and *adherens* junctions form the barrier at the luminal side of the colonic epithelium, while the cell-matrix adhesion molecules form the barrier at the basement membrane between the epithelium and interstitium. Barriers on both sides depend on cell-cell cohesive strength provided by desmosomes (67) as depicted schematically in Figure 6. Inadequacies in any of these barrier-related proteins compromise tissue integrity and likely contribute to interstitial infiltration by bacteria, bacterial toxins and food allergens – all of which are capable of provoking an inflammatory response (68). Not surprisingly, proteins known to have an anti-inflammatory role in the colon were concomitantly up-regulated (48-51).

**Figure 6.**
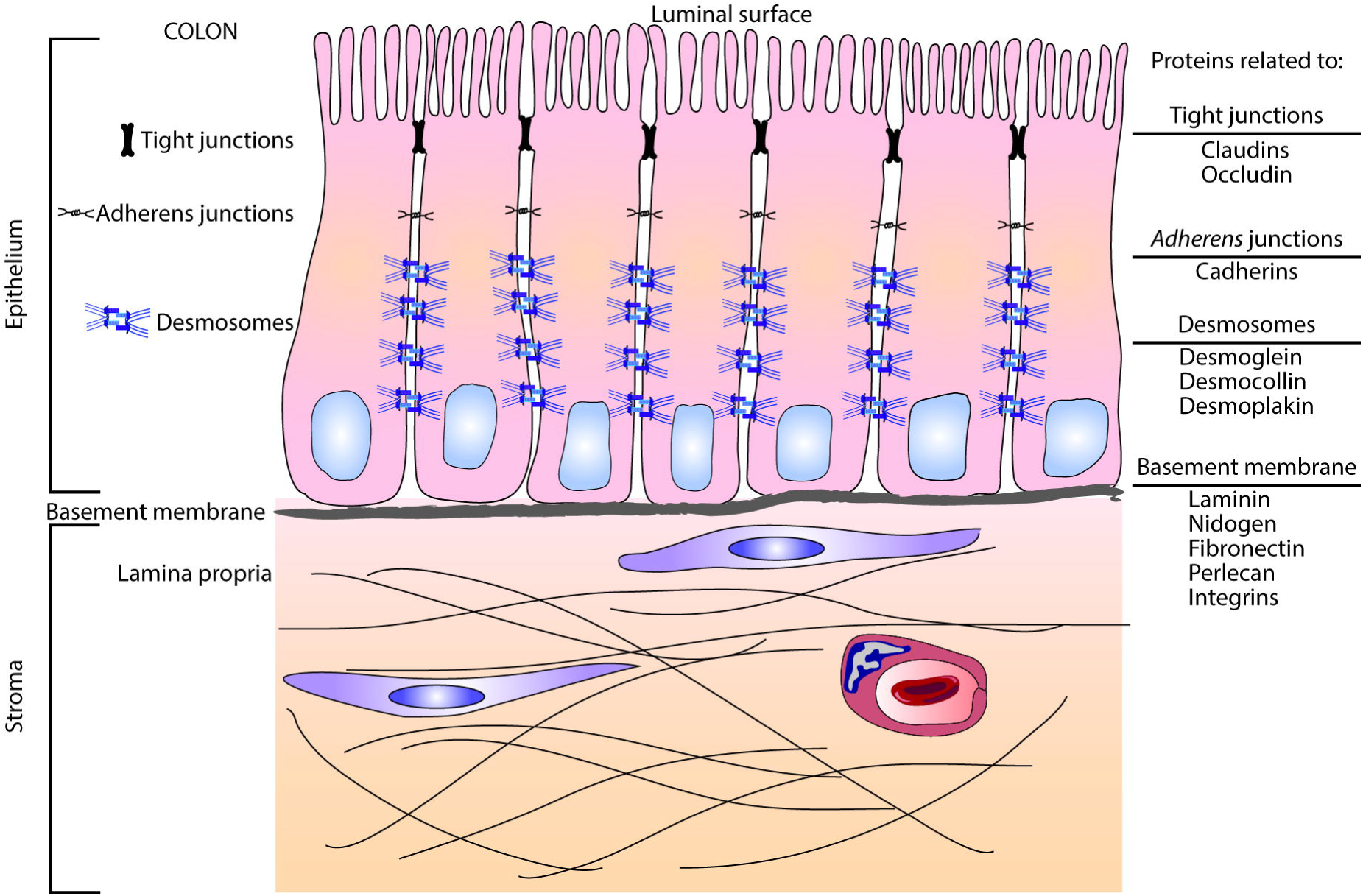
Schematic representation of proteins and protein structures involved in formation of the barrier in the colonic mucosa.

That all of these critical structures are up-regulated with calcium concentrations that are higher than levels needed to promote differentiation, *per se*, attests to the importance of having an adequate calcium-intake level throughout life. Unfortunately, the Western-style diet is deficient in the level of calcium provided to most individuals. This has been well-documented in studies from North America, Europe and Australia (69-71). Additionally, it has been shown that even a substantial percentage of individuals on a mostly plant-based diet do not achieve an adequate calcium intake (72). Calcium is not the only mineral in which there is wide-spread deficiency. Approximately 50% of the US population does not meet the US recommended daily allowance for dietary intake of magnesium (73). While many of the other minerals represented in Aquamin are not routinely tested for and have no daily recommended intake levels, it may be assumed that individuals who do not achieve adequate calcium and magnesium levels would also be deficient in other elements that are nutritionally associated with calcium. Whether or not routine dietary mineral supplementation would be an effective way to mitigate some of the age-related chronic diseases remains to be seen in controlled clinical trials. Such studies are in progress in our laboratory.

In summary, the studies described here demonstrate that colonoids obtained from histologically-normal colon tissue express gross and histological features of differentiation in the presence of a low ambient level of extracellular calcium. In this respect, the normal tissue colonoids are different from the previously-studied adenoma colonoids (38), which demonstrated little evidence of differentiation under low-calcium conditions, but differentiated in response to calcium supplementation. Taken together, these findings suggest that reduced sensitivity to low ambient calcium is an inherent feature of the transformation process in the colonic epithelium. In addition, the present findings show that even in colonoids that are already differentiated based on morphological criteria, intervention with calcium up-regulates proteins that contribute to adhesive function, barrier formation and tissue integrity. To the extent that dietary mineral supplementation can improve barrier formation and tissue integrity in the gastrointestinal tract, it should help reduce chronic inflammation and, ultimately, mitigate some of the age-related conditions that result from chronic inflammation.

## Acknowledgements

This study was supported by National Institutes of Health grants CA181855 and CA201782 (JV) and MCubed (MNA). We thank Marigot LTD (Cork, Ireland) for providing Aquamin^®^ as a gift. We thank the Proteomic Core for help with proteomic data acquisition; the Histology and Immunohistology laboratory (University of Michigan Comprehensive Cancer Center) for immunostaining; the Microscopy and Imaging Laboratory for help with scanning electron microscopy and transmission electron microscopy and the Slide-Scanning Services of the Pathology Department for assistance with slide scanning and morphometric analysis.

## SUPPLEMENTAL MATERIAL

**Supplement Table 1.** Characteristics of subjects providing tissue

**Supplement Table 2.** Antibody characteristics

**Supplement Table 3.** Unbiased analysis of up-regulated proteins

**Supplement Table 4.** Unbiased analysis of down-regulated proteins

**Supplement Figure 1.**
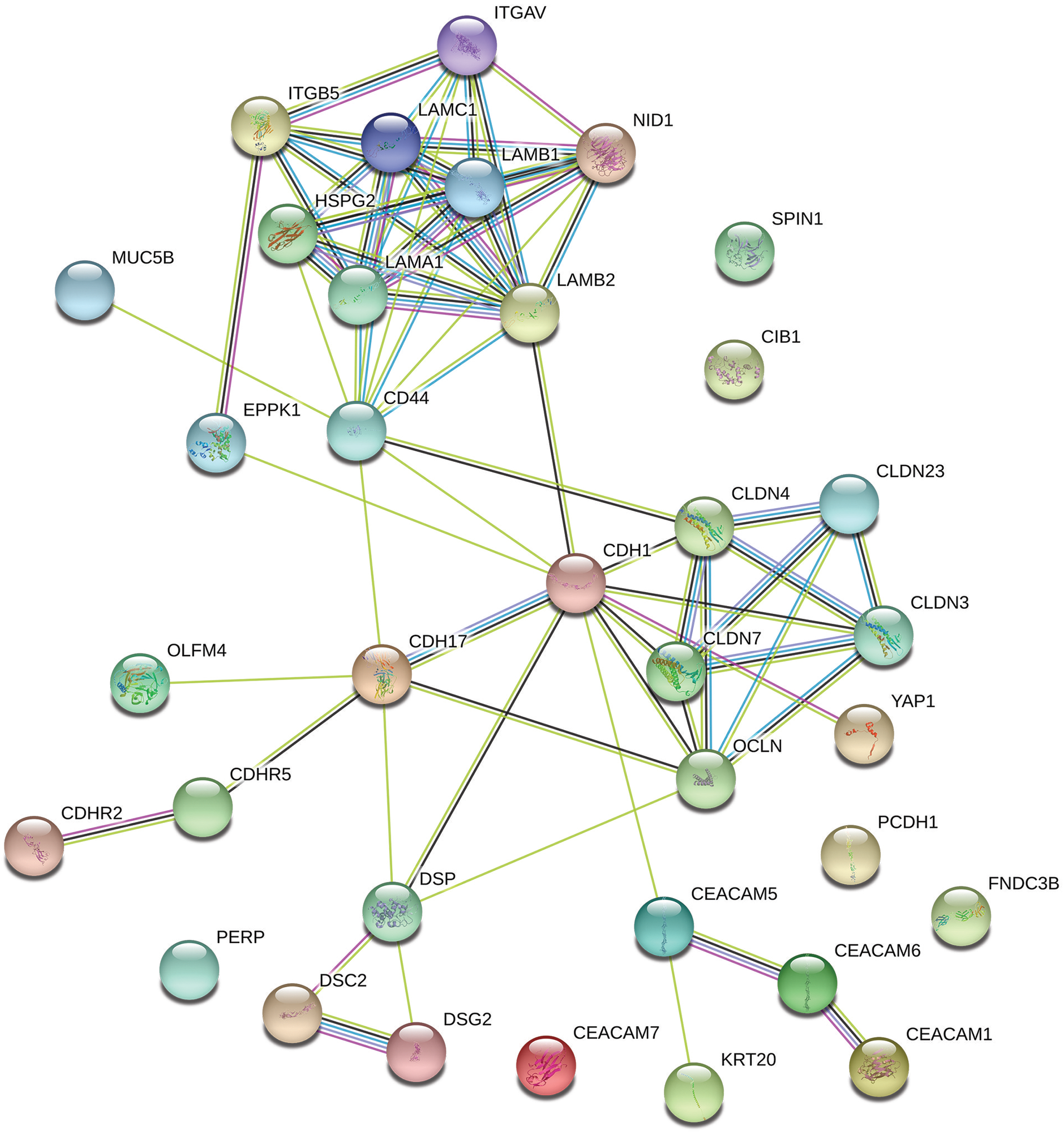
Protein-Protein interactions – String-database.

