## Supplement Table 1 for "Calcium-Induced Differentiation in Normal Human Colonoid Cultures: Cell-cell / cell-matrix adhesion, barrier formation and tissue integrity"

**Supplement Table 1. Human Colon Samples (Used in this study).**

| **Subject ID** | **Age** (Years) **/Sex** | **Location** | **Eligibility criteria** |
|---|---|---|---|
| 90 | 31/F | Sigmoid colon | H/o colon polyp, Family h/o CRC (First degree relative) |
| 104 | 58/F | Sigmoid colon | H/o colon polyp, Family h/o CRC (First degree relative) |
| 106 | 50/M | Sigmoid colon | H/o colon polyp |
| 108 | 61/M | Sigmoid colon | H/o colon polyp, Family h/o CRC (First degree relative) |
| 111 | 20/M | Sigmoid colon | Family h/o CRC (First degree relative) |

H/o = History of, CRC = Colorectal Cancer.
