## Supplement Table 2 for "Calcium-Induced Differentiation in Normal Human Colonoid Cultures: Cell-cell / cell-matrix adhesion, barrier formation and tissue integrity"

**Supplement Table 2.** Antibodies used in the study (for IHC assay)

| Antibody | Vendor | Catalog # | Dilution | Incubation Time | Retrieval Method |
|---|---|---|---|---|---|
| Rb Ki67 MaB clone SP6 | Cell Marque | 475 R-16 | 1:250 | 30 min | FLEX TRS High pH (9.01), 20 min |
| Ms CK20 MaB clone Ks20.8 | Dako | M7019 | 1:100 | 60 min | FLEX TRS High pH (9.01), 20 min |
| Rb LI Cadherin (CDH17) MaB clone EPR3996 | AbCam | Ab109190 | 1:250 | 60 min | [a]HIER pH 9.0 |
| Rb Desmoglein 2 PaB | Sigma | HPA004896 | 1:200 | 30 min | [b]HIER pH 6.0 |
| Rb Claudin 23 PaB | AbCam | Ab23355 | 1:100 | 30 min | [a]HIER pH 9.0 |

[a]HIER pH 9: Heat induced epitope retrieval 10 mM Tris HCl/1 mM EDTA buffer pH9

[b]HIER pH 6: Heat induced epitope retrieval Citrate buffer pH6
