## Supplement Table 3 for "Calcium-Induced Differentiation in Normal Human Colonoid Cultures: Cell-cell / cell-matrix adhesion, barrier formation and tissue integrity"

**Supplement Table 3A. Up-regulated proteins (Unbiased approach).**

| Protein | Calcium 1.5mM | Calcium 2.1mM | Calcium 3.0mM | Aquamin (1.5mM Calcium) | Aquamin (2.1mM Calcium) | Aquamin (3.0mM Calcium) |
|---|---|---|---|---|---|---|
| Keratin, type I cytoskeletal 9 | 1.07±0.47 | 1.79±1.73 | *0.59±0.13 | 3.66±4.03 | 9.96±15.39 | 0.98±0.34 |
| Keratin, type II cytoskeletal 1 | 0.94±0.30 | 1.58±1.41 | 0.73±0.34 | 2.79±3.22 | 7.12±8.56 | 0.95±0.12 |
| Keratin, type I cytoskeletal 16 | 0.98±0.14 | 3.88±4.88 | 1.00±0.14 | 5.59±8.08 | 8.11±7.19 | 0.92±0.22 |
| Hornerin | 1.68±1.22 | 1.82±1.28 | 0.76±0.09 | 1.99±0.95 | 4.98±6.49 | 0.94±0.62 |
| Keratin, type II cytoskeletal 6A | 0.92±0.16 | 1.80±1.59 | 0.98±0.07 | 3.95±5.37 | 4.95±5.38 | 1.07±0.25 |
| Keratin, type I cytoskeletal 14 | 0.84±0.25 | 3.33±3.45 | 1.24±0.71 | 1.88±1.89 | 4.36±2.68 | 0.99±0.14 |
| Keratin, type II cytoskeletal 5 | 1.14±0.34 | 1.47±0.60 | 1.08±0.15 | 2.83±3.09 | 4.50±3.04 | 0.98±0.02 |
| Cadherin-17 | *4.28±1.04 | *4.57±1.43 | *5.65±0.69 | *3.89±0.90 | *3.87±0.98 | *4.62±0.25 |
| Vitamin D-binding protein | 2.99±1.28 | 3.29±2.50 | 3.49±2.12 | *2.04±0.49 | 2.27±0.85 | 1.75±0.56 |
| Carbonic anhydrase 1 | 1.70±0.51 | 1.83±0.73 | *1.96±0.17 | 2.16±1.97 | 2.10±1.87 | 1.02±0.29 |
| Lactotransferrin | 3.08±1.81 | 3.09±1.86 | *1.94±0.15 | 2.33±1.10 | 2.74±1.52 | 1.74±0.88 |
| Sulfate transporter | 2.54±1.04 | 2.31±1.02 | *2.20±0.15 | 3.19±2.09 | 2.96±2.14 | *1.91±0.20 |
| Zinc transporter ZIP4 | 2.28±1.51 | 2.00±1.04 | 1.51±0.86 | 2.30±0.90 | 2.45±1.56 | 1.68±0.78 |
| Natural resistance-associated macrophage protein 2 | 2.45±1.44 | 2.06±0.70 | *1.65±0.20 | *2.48±0.80 | 2.58±1.49 | *2.27±0.13 |
| Alpha-2-HS-glycoprotein | 2.57±1.07 | 2.76±1.19 | 2.82±0.86 | *2.10±0.6 | 2.42±1.36 | 1.45±0.42 |
| Aminopeptidase N | *2.31±0.56 | *2.05±0.52 | *2.40±0.32 | 2.46±0.93 | 2.28±1.27 | 1.68±0.37 |
| Prostate stem cell antigen | *2.50±0.64 | *2.20±0.72 | *2.55±0.15 | 2.27±0.97 | 2.08±1.41 | 1.59±0.94 |
| Xaa-Pro aminopeptidase 2 | *1.91±0.55 | 1.84±0.63 | 1.47±0.26 | 2.38±0.96 | 2.40±1.09 | 1.64±0.36 |
| Intestinal-type alkaline phosphatase | *2.33±0.76 | *2.16±0.54 | *2.29±0.26 | *2.38±0.56 | 2.35±0.98 | 1.82±0.70 |
| Alpha-2-macroglobulin | *2.55±0.39 | *2.48±0.78 | *2.80±0.15 | *2.18±0.14 | *2.79±0.40 | *2.70±0.005 |
| Meprin A subunit alpha | *2.07±0.65 | *2.01±0.53 | *2.13±0.17 | 2.30±0.83 | 2.01±1.14 | 1.51±0.76 |
| Chloride anion exchanger | *1.94±0.47 | *1.81±0.42 | *1.87±0.11 | 2.23±0.95 | 2.05±1.00 | 1.56±0.51 |
| CEACAM7 | *2.05±0.52 | *1.92±0.46 | *1.94±0.04 | 2.26±0.88 | 2.13±0.93 | 1.59±0.40 |
| Desmoglein-2 | *2.52±0.42 | *2.54±0.61 | *3.15±0.06 | *2.40±0.52 | *2.37±0.57 | *2.70±0.07 |

| | | | | | | |
|---|---|---|---|---|---|---|
| Complement C3 | *2.58±0.36 | *2.41±0.65 | *2.71±0.56 | *2.00±0.28 | *2.38±0.71 | *1.89±0.11 |
| Trefoil factor 2 | *1.83±0.33 | *1.91±0.25 | *1.93±0.28 | 1.95±0.94 | 1.75±0.84 | 1.45±0.64 |
| Protocadherin-1 | *2.21±0.43 | *2.48±0.52 | 2.37±0.84 | *1.90±0.38 | *2.13±0.18 | *2.17±0.25 |
| 15-hydroxyprostaglandin dehydrogenase [NAD(+)] | 1.90±0.61 | *1.91±0.53 | *2.03±0.27 | 1.91±0.71 | 1.96±0.83 | 1.42±0.22 |
| Solute carrier family 15 member 1 | 1.91±0.57 | 1.68±0.46 | *1.95±0.14 | 1.95±0.71 | 1.85±0.89 | 1.36±0.34 |
| Calcium-activated chloride channel regulator 4 | *1.77±0.43 | *1.75±0.31 | *2.07±0.13 | 1.87±0.79 | 2.00±0.82 | 1.55±0.63 |
| Hydroxymethylglutaryl-CoA synthase, mitochondrial | *1.78±0.37 | *1.82±0.39 | *1.88±0.11 | *1.88±0.38 | *1.99±0.16 | *2.07±0.13 |
| Pantetheinase | 1.46±0.35 | *1.39±0.24 | *1.76±0.14 | 1.40±0.38 | 1.27±0.50 | 1.17±0.15 |

Values represent average fold change from 3 colonoids as compared to control (0.25 mM calcium) ±SD. These proteins were up-regulated in all 3 colonoids at 1.8-fold change and were common in all three colonoids based on a maximum up-regulation in at least one condition. Some of the common up-regulated proteins have presented in table 1 as part of the differentiation-related panel. *Represents significance as compared to the control at $p<0.05$.

**Supplement Table 3B. Top pathways associated with Up-regulated proteins (Reactome v66)**

| Pathway name | Entities pValue |
|---|---|
| Formation of the cornified envelope | 0.0000001 |
| Keratinization | 0.000002 |
| Post-translational modification: synthesis of GPI-anchored proteins | 0.00001 |
| SLC transporter disorders | 0.00004 |
| Multifunctional anion exchangers | 0.0003 |
| Neutrophil degranulation | 0.0004 |
| Type I hemidesmosome assembly | 0.0005 |
| SLC-mediated transmembrane transport | 0.001 |
| Disorders of transmembrane transporters | 0.001 |
| Transport of small molecules | 0.001 |
| Cell junction organization | 0.002 |
| Metal ion SLC transporters | 0.003 |
| Defective SLC39A4 causes acrodermatitis enteropathica, zinc-deficiency type (AEZ) | 0.003 |
| Defective SLC11A2 causes hypochromic microcytic anemia, with iron overload 1 (AHMIO1) | 0.003 |
| Defective SLC26A2 causes chondrodysplasias | 0.003 |
| Defective SLC26A3 causes congenital secretory chloride diarrhea 1 (DIAR1) | 0.003 |
| Transport of inorganic cations/anions and amino acids/oligopeptides | 0.004 |
| Cell-Cell communication | 0.006 |
| Proton/oligopeptide cotransporters | 0.011 |
| Biosynthesis of D-series resolvins | 0.011 |
| Alternative complement activation | 0.014 |
| Biosynthesis of E-series 18(S)-resolvins | 0.014 |
| Mtb iron assimilation by chelation | 0.017 |
| Transport and synthesis of PAPS | 0.017 |
| Metal sequestration by antimicrobial proteins | 0.017 |
| Synthesis of Lipoxins (LX) | 0.017 |
| Biosynthesis of EPA-derived SPMs | 0.017 |
| Activation of C3 and C5 | 0.020 |

| | |
|---|---|
| Erythrocytes take up oxygen and release carbon dioxide | 0.023 |
| Synthesis of Ketone Bodies | 0.023 |
| HDL assembly | 0.023 |
| Transport of bile salts and organic acids, metal ions and amine compounds | 0.026 |
| Developmental Biology | 0.028 |
| Zinc influx into cells by the SLC39 gene family | 0.028 |
| Ketone body metabolism | 0.028 |
| Vitamin D (calciferol) metabolism | 0.031 |
| Apoptotic cleavage of cell adhesion proteins | 0.031 |
| Reversible hydration of carbon dioxide | 0.034 |
| O2/CO2 exchange in erythrocytes | 0.034 |
| Erythrocytes take up carbon dioxide and release oxygen | 0.034 |
| Post-translational protein phosphorylation | 0.038 |
| Synthesis of Prostaglandins (PG) and Thromboxanes (TX) | 0.042 |
| Innate Immune System | 0.048 |
| Vitamin B5 (pantothenate) metabolism | 0.048 |
| Zinc transporters | 0.048 |
| Metabolism of Angiotensinogen to Angiotensins | 0.048 |
| Biosynthesis of DHA-derived SPMs | 0.048 |
| Regulation of Insulin-like Growth Factor transport and uptake by Insulin-like Growth Factor Binding Proteins | 0.050 |
