## Supplement Table 4 for "Calcium-Induced Differentiation in Normal Human Colonoid Cultures: Cell-cell / cell-matrix adhesion, barrier formation and tissue integrity"

**Supplement Table 4A. Down-regulated proteins (Unbiased approach).**

| Protein | Calcium 1.5mM | Calcium 2.1mM | Calcium 3.0mM | Aquamin (1.5mM Calcium) | Aquamin (2.1mM Calcium) | Aquamin (3.0mM Calcium) |
|---|---|---|---|---|---|---|
| Coiled-coil domain-containing protein 86 | *0.377±0.266 | *0.317±0.247 | *0.255±0.143 | *0.462±0.157 | *0.341±0.174 | *0.279±0.144 |
| Caveolae-associated protein 1 | *0.551±0.174 | 0.807±0.684 | *0.445±0.074 | *0.455±0.139 | *0.386±0.247 | *0.451±0.019 |
| Proliferating cell nuclear antigen | 0.677±0.279 | 0.796±0.435 | *0.485±0.146 | *0.608±0.134 | *0.556±0.099 | *0.605±0.069 |
| Inosine-5'-monophosphate dehydrogenase 2 | *0.598±0.209 | *0.614±0.17 | *0.453±0.040 | *0.523±0.081 | *0.526±0.126 | *0.487±0.076 |
| Importin subunit alpha-1 | 0.785±0.415 | 0.922±0.707 | *0.515±0.054 | *0.674±0.159 | *0.566±0.150 | 0.621±0.150 |
| 60S ribosomal protein L36 | *0.538±0.054 | *0.551±0.132 | *0.559±0.087 | *0.769±0.080 | *0.564±0.073 | *0.654±0.017 |
| RNA-binding protein with serine-rich domain 1 | *0.708±0.057 | *0.643±0.12 | *0.476±0.060 | *0.781±0.124 | 0.742±0.190 | 0.819±0.094 |
| Nucleophosmin | *0.590±0.104 | 0.714±0.255 | 0.607±0.161 | *0.603±0.081 | *0.541±0.097 | *0.590±0.076 |

Values represent average fold change from 3 colonoids as compared to control (0.25 mM calcium) ±SD. These proteins were down-regulated in all 3 colonoids at 1.8-fold change and were common in all three colonoids based on a maximum down-regulation in at least one condition. *Represents significance as compared to the control at $p<0.05$.

**Supplement Table 4B. Pathways associated with Down-regulated proteins (Reactome v66)**

| Pathway name | Entities pValue |
|---|---|
| TP53 Regulates Transcription of Cell Cycle Genes | 0.0005 |
| Chromosome Maintenance | 0.0018 |
| Nonsense-Mediated Decay (NMD) | 0.003 |
| Nonsense Mediated Decay (NMD) enhanced by the Exon Junction Complex (EJC) | 0.003 |
| Infectious disease | 0.003 |
| TFAP2A acts as a transcriptional repressor during retinoic acid induced cell differentiation | 0.004 |
| Sensing of DNA Double Strand Breaks | 0.004 |
| DNA Double-Strand Break Repair | 0.005 |
| Influenza Infection | 0.006 |
| Regulation of expression of SLITs and ROBOs | 0.006 |
| SUMO E3 ligases SUMOylate target proteins | 0.006 |
| SUMOylation | 0.007 |
| Removal of the Flap Intermediate from the C-strand | 0.007 |
| Processive synthesis on the C-strand of the telomere | 0.008 |
| Purine ribonucleoside monophosphate biosynthesis | 0.009 |
| Signaling by ROBO receptors | 0.010 |
| Removal of the Flap Intermediate | 0.010 |
| Polymerase switching | 0.010 |
| Leading Strand Synthesis | 0.010 |
| Polymerase switching on the C-strand of the telomere | 0.010 |
| Mismatch repair (MMR) directed by MSH2:MSH3 (MutSbeta) | 0.010 |
| Mismatch repair (MMR) directed by MSH2:MSH6 (MutSalpha) | 0.010 |
| Processive synthesis on the lagging strand | 0.011 |
| Mismatch Repair | 0.011 |
| Nucleobase biosynthesis | 0.011 |
| Translesion synthesis by REV1 | 0.011 |
| Translesion synthesis by POLI | 0.012 |
| Translesion synthesis by POLK | 0.012 |
| TP53 Regulates Transcription of Genes Involved in G2 Cell Cycle Arrest | 0.013 |
| Gene expression (Transcription) | 0.013 |
| Translesion Synthesis by POLH | 0.014 |
| Transcription of E2F targets under negative control by DREAM complex | 0.014 |
| Lagging Strand Synthesis | 0.014 |

| | |
|---|---|
| PCNA-Dependent Long Patch Base Excision Repair | 0.015 |
| TP53 regulates transcription of additional cell cycle genes | 0.015 |
| Telomere C-strand (Lagging Strand) Synthesis | 0.017 |
| Gap-filling DNA repair synthesis and ligation in GG-NER | 0.018 |
| Resolution of AP sites via the multiple-nucleotide patch replacement pathway | 0.018 |
| DNA Repair | 0.018 |
| G0 and Early G1 | 0.019 |
| Activation of E2F1 target genes at G1/S | 0.020 |
| G1/S-Specific Transcription | 0.020 |
| Extension of Telomeres | 0.021 |
| Recognition of DNA damage by PCNA-containing replication complex | 0.022 |
| RNA Polymerase I Transcription Termination | 0.022 |
| Termination of translesion DNA synthesis | 0.023 |
| DNA strand elongation | 0.023 |
| Nuclear import of Rev protein | 0.026 |
| Transcriptional regulation by the AP-2 (TFAP2) family of transcription factors | 0.026 |
| Transcriptional Regulation by TP53 | 0.027 |
| Translesion synthesis by Y family DNA polymerases bypasses lesions on DNA template | 0.028 |
| Resolution of Abasic Sites (AP sites) | 0.028 |
| Base Excision Repair | 0.028 |
| Interactions of Rev with host cellular proteins | 0.028 |
| Dual Incision in GG-NER | 0.029 |
| NS1 Mediated Effects on Host Pathways | 0.031 |
| SUMOylation of transcription cofactors | 0.031 |
| Host Interactions with Influenza Factors | 0.034 |
| SUMOylation of DNA replication proteins | 0.034 |
| DNA Damage Bypass | 0.035 |
| Deposition of new CENPA-containing nucleosomes at the centromere | 0.038 |
| Nucleosome assembly | 0.038 |
| mRNA 3'-end processing | 0.040 |
| E3 ubiquitin ligases ubiquitinate target proteins | 0.042 |
| DNA Double Strand Break Response | 0.042 |
| Disease | 0.043 |
| Telomere Maintenance | 0.045 |
| Gap-filling DNA repair synthesis and ligation in TC-NER | 0.045 |
| Dual incision in TC-NER | 0.046 |

| | |
|---|---|
| RNA Polymerase II Transcription Termination | 0.047 |
| Cleavage of Growing Transcript in the Termination Region | 0.047 |
| HDR through Homologous Recombination (HRR) | 0.047 |
